# Predicted structure of the 89-kDa invasion-tip protein of *Holospora obtusa*, an endonuclear symbiont of *Paramecium caudatum*, suggests a pH-dependent conformational change

**DOI:** 10.64898/2026.10.08.757700

**Authors:** Yasuhiro Takenaka, Toshiyuki Murai, Toru Nakatsu, Yasushi Shigeri, Masahiro Fujishima

## Abstract

The endonuclear symbiont *Holospora obtusa* invades the macronucleus of *Paramecium caudatum* using an infectious-form invasion-tip containing a major 89-kDa periplasmic protein implicated in escape from the host digestive vacuole and macronuclear invasion. Sequence-based analyses placed this protein within a *Holospora*-specific family and predicted an N-terminal membrane-anchoring module and a long *α*-helical C-terminal region. AlphaFold2 modeling followed by molecular dynamics indicated that the C-terminal region forms an antiparallel, spectrin-repeat-like helical solenoid containing a localized three-helix coiled-coil. The protein exhibits a large pH-dependent charge difference, becoming 55 units more positive at pH 4 than at pH 7. Independent simulations at these pH values differed mainly in the relative orientation of the N-terminal core and C-terminal rod, described by DynDom as an approximately 35° rigid-body rotation about a hinge centered on Gly291. However, none of the nearby glutamates (Glu273, Glu275, and Glu302) formed a pH-dependent salt bridge, and no specific protonation-dependent interaction responsible for the conformational difference was identified. We nonetheless propose, as a testable hypothesis, that acidification of the digestive vacuole may promote this reorientation, thereby repositioning the N-terminal membrane-anchoring module at the bacterial surface as an early step toward escape from the vacuole.

## 1. Introduction

*Holospora obtusa* is a Gram-negative alphaproteobacterial endonuclear symbiont of the ciliate *Paramecium caudatum*. A highly specialized infection strategy has evolved that enables it to invade the host macronucleus while surviving entirely within the intracellular environment [1–6]. Unlike free-living bacteria, *Holospora* species have undergone substantial genome reduction [2,3,7] and cannot be cultivated outside their host cells, which indicates a high degree of adaptation to intracellular life. During their life cycle, they show two morphologically different forms: a reproductive short form (RF, 1.5–2 µm in length) and an infectious long form (IF, 10–15 µm in length) [8–10]. When the host is growing by binary fission, the RF cell proliferates by binary fission within the macronucleus. If the host cell starves, the RF ceases the binary fission and differentiates into the IF cell via intermediate forms. The IF cell forms a characteristic structure consisting of two regions: one containing cytoplasm and the other a large periplasmic region with an electron-lucent invasion-tip [1,11–15]. Infection can be initiated synchronously by mixing isolated IF cells from host cell homogenates [16] with uninfected host cells. The *Holospora–Paramecium* system therefore provides an exceptional experimental model for investigating the earliest molecular events that enable endosymbiotic bacteria to recognize environmental cues, escape from a host digestive compartment, and invade its specific intracellular niche.

Shortly after uptake by the host, IF cells are enclosed within newly formed digestive vacuoles (DVs). DVs of *Paramecium* species have been classified into four stages [16–18]. The DV-I is a young vacuole produced by pinching off from a cytopharynx. The DV-I fuses with acidosomes and transforms into a contracted and acidified DV-II (inside pH < 3), which then fuses with primary lysosomes and differentiates to DV-III. DV-IV is the final stage of the DV before defecation and lacks acid phosphatase activity. In the DV-I, the IF cells rapidly undergo a series of coordinated morphological and physiological changes that transform them into activated forms (AFs), and this transformation is inducible if IFs are treated with a low pH buffer (pH 4) for 15 min [19]. The AF cell escapes only from the DV-I, migrates through the host cytoplasm, and penetrates the macronuclear envelope by leading with the invasion-tip. Escape occurs only during the brief period in which the DV lumen is undergoing acidification. Bacterial escape is completely blocked once the vacuole matures to the more acidic DV-II stage. Furthermore, inhibition of vacuolar acidification by the V-type H^+^-ATPase inhibitors concanamycin A or concanamycin B abolishes both escape from the DV and subsequent nuclear infection [20]. These observations have established transient acidification in the DV-I as the essential environmenatl signal that initiates the infection process. However, the bacterial molecule that senses this pH change and converts it into the structural changes required for membrane engagement and host-cell invasion is unknown.

To identify the bacterial factor responsible for this acid-triggered activation process, we previously generated a monoclonal antibody against a major 89-kDa protein localized specifically within the lumen of the invasion-tip and subsequently identified its full-length gene [14]. Sequence analysis revealed a hydrophobic N-terminal segment, annotated at the time as a signal peptide, followed immediately by two putative actin-binding motifs (WH2 and FH2). This suggests that this protein might directly connect environmental sensing with interactions between the bacterial surface and the host cytoskeleton. Immunofluorescence microscopy further demonstrated that, although the protein is initially distributed uniformly throughout the invasion-tip lumen in freshly isolated IFs, its N-terminal region rapidly becomes exposed on the outer surface of the bacterial outer membrane immediately after bacterial uptake into host DVs. At the same stage, transmission electron microscopy revealed numerous filamentous structures bridging the bacterial surface and the DV membrane [14]. Following escape into the host cytoplasm, host actin accumulated around the exposed N-terminal region [16,21], and similar actin-associated structures remained at the site where the bacterium penetrated the macronuclear envelope [16]. Together, these observations identified the 89-kDa protein as a central component throughout the entire infection process, including escape from the DV, migration through the host cytoplasm, and penetration of the target nucleus.

Nonetheless, one fundamental question has remained unanswered. Although the N-terminal region of the 89-kDa protein clearly becomes exposed during infection, the molecular mechanism by which acidification of the DV triggers this dramatic structural rearrangement has not been determined. Specifically, there has been no explanation of how exposure to the acidic environment of the DV-I converts a protein that is initially confined within the invasion-tip lumen into one capable of interacting with the bacterial outer membrane and, subsequently, with the host actin cytoskeleton. The molecular basis linking DV acidification to activation of the invasion machinery has thus remained unresolved for nearly two decades.

To elucidate the molecular mechanism underlying acid-triggered activation of *H. obtusa* infection, we investigated the three-dimensional structure and pH-dependent conformational dynamics of the 89-kDa periplasmic protein using AlphaFold2-based structural prediction and molecular dynamics simulations. Our objective was to determine whether pH-dependent conformational changes in this protein could provide a structural basis for activation of the bacterial invasion machinery. In addition, to determine whether proteins with similar amino acid sequences have been identified in other organisms, we also performed an updated homology search of the 89-kDa protein sequence using current sequence databases.

## 2. Materials and Methods

### 2.1. Phylogenetic Analyses

A BLASTP search against the nr database (accessed July 2026; default parameters) was used to assess similarity to proteins inside and outside the genus *Holospora*. Multiple alignment was performed using MUSCLE with default parameters in MacVector version 18.8.3, using a log-expectation profile. The best-fit substitution model was selected by model testing in MEGA version 12.1.2 [22], and the JTT+G+I model, which showed the lowest Bayesian information criterion score, was used for tree construction.

### 2.2. Computational Modeling and Molecular Dynamics Simulations

The three-dimensional structure of the 89-kDa periplasmic protein (BAE80122.1; 750 residues) was predicted using ColabFold [23] with AlphaFold2 [24] default parameters (run October 2025). The top-ranked model was selected for subsequent simulations.

pH-dependent protonation states were assigned using PDB2PQR version 3.7.1 with PROPKA [25,26] at pH 4.0 and pH 7.0. At pH 4.0, seven histidine residues (His96, His133, His364, His542, His588, His615, His735) were treated as doubly protonated (HIP), eight aspartate residues (Asp125, Asp225, Asp411, Asp444, Asp551, Asp585, Asp609, Asp611) were assigned as neutral protonated forms (ASH), and 39 glutamate residues were treated as neutral protonated forms (GLH), resulting in a net charge shift of +55 elementary charges relative to the pH 7.0 state.

Each protonation variant was parameterized with the AMBER ff14SB force field [27] and solvated in an explicit TIP3P water box [28] with a minimum padding of 10 Å between the protein surface and the box boundary. Counter-ions (Cl^−^) were added to electro-neutralize the system (59 Cl^−^ for pH 4.0; 4 Cl^−^ for pH 7.0). MD simulations were performed using OpenMM version 8.3.1 [29]. Each system was energy-minimized (up to 10,000 iterations), followed by stepwise heating from 0 to 300 K in increments of 50 K (5,000 steps per increment at 1 fs timestep). Equilibration was then carried out for 100 ps at 300 K with a 2 fs timestep. Production runs of 10 ns were performed under NPT conditions (300 K, 1 atm) using a LangevinMiddle integrator (friction coefficient 1 ps^−1^, timestep 2 fs) with a Monte Carlo barostat (coupling every 25 steps). Particle mesh Ewald summation [30] was used for long-range electrostatics (cutoff 1.0 nm, Ewald error tolerance 0.0005), with hydrogen bond lengths constrained and rigid water geometry enforced. Coordinates were saved every 10 ps, yielding 1,000 frames per simulation.

### 2.3. Secondary-structure, Coiled-coil and Membrane-topology Prediction

The amino-acid sequence of the 89-kDa periplasmic protein of *H. obtusa* (750 aa) was analyzed with a panel of sequence-based predictors using the MPI Bioinformatics Toolkit (Tübingen, Germany) [31]. Consensus secondary structure was obtained with Quick2D, which integrates PSIPRED [32], SPIDER3 [33], PSSpred, DeepCNF/RaptorX-Property [34] and NetSurfP-2.0 [35]. Coiled-coil propensity was obtained with COILS and PCOILS [36,37] (window size 28) and MARCOIL [38]; transmembrane segments with TMHMM 2.0 [39]. Intrinsic disorder was obtained with NetSurfP-2.0, DISOPRED3 [40], SPOT-Disorder and IUPred. Coiled-coil propensity was additionally evaluated with DeepCoil2 [41], reporting residues with a coiled-coil probability ≥ 0.5. A predicted signal peptide (PSG, GvH), transmembrane segment (ALOM), membrane topology (MTOP), coiled-coil region (Lupas COIL) and subcellular localization (k-NN) were obtained with PSORT II [42]. All servers were run with default parameters (accessed June 2026).

Structure-based analyses were performed on the final frame of the pH 7 molecular-dynamics trajectory. Secondary structure was assigned from C*α* coordinates using the P-SEA method [43], giving a three-state (helix/strand/coil) assignment equivalent to DSSP [44]. Knobs-into-holes packing was identified with SOCKET2 (SOCKET v3.02) [45] using a packing-cutoff of 7.0 Å on the heavy-atom model. C*α*–C*α* contact maps (contacts defined at < 12 Å) and an internal structural-repeat scan of the C-terminal rod (residues 198–720), based on sliding-window superposition and contact-map autocorrelation, were computed in-house with Python 3 and NumPy to detect tandem antiparallel *α*-helical modules.

### 2.4. Homology Searches

Remote homology was assessed with HHpred (MPI Bioinformatics Toolkit; accessed May 2026) using the amino-acid sequence of BAE80122.1 as the query. Structural similarity was assessed with Foldseek (databases AFDB50 and MGnify-ESM Atlas), using the final frame of the pH 7 molecular-dynamics trajectory as the query structure. The ten highest-scoring hits per structural class or database are listed in Tables S1 and S2, respectively.

### 2.5. Structural Analysis

The final frame of each 10-ns trajectory was extracted as the representative equilibrated structure (Files S1 and S2). Per-residue C*α* RMSF was calculated from the last 500 frames (final 5 ns) of each trajectory. Structures were superimposed on the C-terminal rod or on the N-terminal core using Biopy-thon’s Superimposer module prior to C*α* displacement calculations. Domain boundaries and hinge regions were identified using DynDom [46] (http://dyndom.cmp.uea.ac.uk/dyndom/). Structural figures were prepared with UCSF ChimeraX version 1.10.1.

## 3. Results

### 3.1. Phylogenetic analysis of the 89-kDa periplasmic protein from Holospora obtusa

To examine the evolutionary relationship of the 89-kDa periplasmic protein from *Holospora obtusa* BAE80122.1, we first searched for homologous amino acid sequences and performed multiple sequence alignment. A BLASTP search of BAE80122.1 against the nr database using default parameters showed no hits to proteins outside the genus *Holospora*, so subsequent analyses were restricted to *Holospora* homologs. The alignment revealed several conserved amino acid residues or short conserved regions among the *Holospora* proteins, suggesting that these residues may contribute to structural or functional features shared by this protein family (Figure 1A).

**Figure 1.**
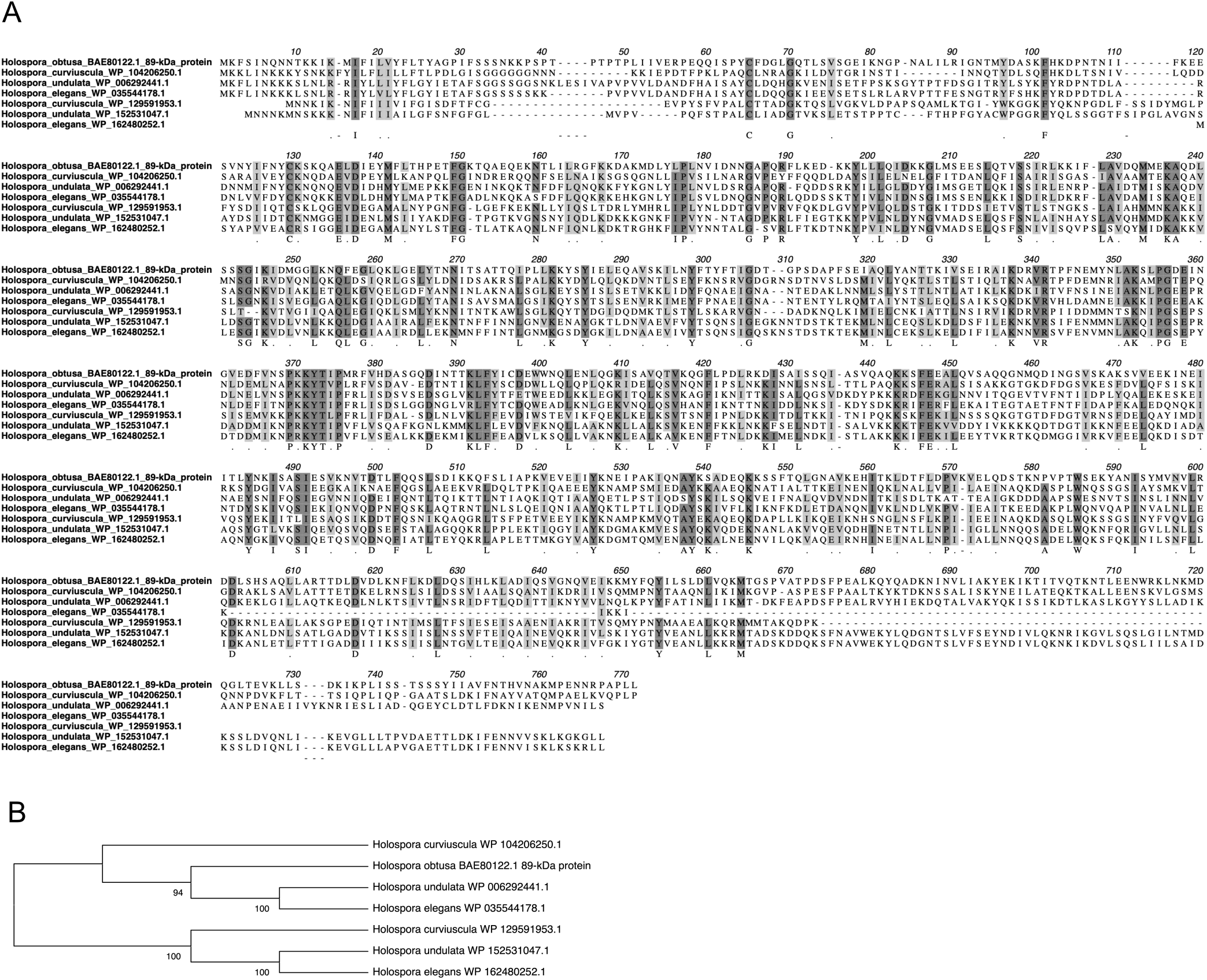
Phylogeny of the 89-kDa periplasmic invasion-tip protein from *Holospora obtusa*. (**A**) Multiple amino acid sequence alignment of the 89-kDa periplasmic protein and *Holospora* homologs. Darkly and lightly shaded text represents at least 80% identical and similar amino acid residues, respectively. NCBI accession numbers are indicated with species names. (**B**) The ML consensus tree obtained from bootstrap analysis with 1,000 replications of the 89-kDa periplasmic protein based on amino acid sequence alignments shown in (**A**). Bootstrap values > 60% are given to the left of selected nodes.

We next constructed a maximum-likelihood phylogenetic tree based on the amino acid alignment. Bootstrap analysis with 1,000 replications showed that the *Holospora* homologs were separated into two major clades (Figure 1B). The 89-kDa periplasmic protein from *H. obtusa* BAE80122.1 clustered with one group of *Holospora* proteins, together with homologs from *H. curviuscula, H. undulata*, and *H. elegans*. A second clade contained additional homologous proteins from these species, suggesting that multiple related proteins may be present within individual *Holospora* species. These results indicate that the 89-kDa periplasmic protein belongs to a conserved *Holospora*-specific protein family. The separation of the homologs into two clades suggests possible diversification of this protein family, potentially through gene duplication, although this possibility requires further genomic and phylogenetic analysis. Notably, a corresponding type-II homolog has not yet been identified for *H. obtusa*, which may reflect incomplete sequence availability or, alternatively, true absence of this paralog in the currently available dataset. Further analysis of the conserved residues identified in the alignment, particularly in relation to the domain organization and pH-dependent structural rearrangement described below, may help to identify residues that are important for the structural behavior of BAE80122.1.

To characterize the protein in the absence of a close homologue, we applied a panel of primary-sequence-based prediction tools. Consensus secondary-structure prediction (PSIPRED, SPIDER3, PSSpred, DeepCNF and NetSurfP-2.0) divided the protein into a short N-terminal module (residues 1–197) of mixed *α*/β character with low-complexity, partly disordered segments and only ∼29% helix, and a long C-terminal region (residues 198–747) predicted to be almost entirely *α*-helical (∼82% helix; Figure S1A). The N-terminal module contained a single strongly hydrophobic segment predicted as a transmembrane/type-II signal-anchor helix (residues 15–31; TMHMM and PSORT II/ALOM) rather than a cleavable signal peptide. This segment was originally annotated as a signal peptide [14]; the two assignments differ only in whether a cleavage site is present, and current predictors favor the non-cleaved form. Coiled-coil prediction assigned propensity to a single short locus near residues 405–437, on which DeepCoil2, COILS, MARCOIL and the Lupas COIL algorithm (PSORT II) agreed; only PCOILS, a known over-predictor on continuous *α*-helical regions, predicted a broader region as ‘coiled-coil’ (Figure S1B). These predictions indicated that the C-terminal two-thirds of the protein forms an elongated *α*-helical region with a single coiled-coil locus. However, we could not distinguish a continuous coiled-coil from an antiparallel helical-repeat, so we structurally examined the region using a three-dimensional model (below).

### 3.2. Molecular modeling and structure-based analysis of the C-terminal rod

To characterize the higher-order architecture of the 89-kDa periplasmic protein, we examined computational models of the protein. The structure of BAE80122.1 (750 residues) was predicted with AlphaFold2, and the ribbon model was colored by the per-residue confidence score (pLDDT) (Figure S2A). Most of the protein body is predicted with high confidence (pLDDT > 90, blue) and forms an elongated array of *α*-helices. In contrast, the N-terminal region (residues 1–30) shows markedly low pLDDT scores (< 50, yellow), so the conformation of this segment in the model is not well determined. The putative transmembrane segment (residues 15–31) lies within this low-confidence region. Ionization states were then assigned with PDB2PQR/PROPKA, using the same procedure at pH 7 and at pH 4 (Figure S2B). Of the 55 residues predicted to differ between the two conditions, 39 Glu and eight Asp residues are neutral at pH 4 and negatively charged at pH 7, seven His residues carry a positive charge at pH 4 and are neutral at pH 7, and one lysine (Lys155) is protonated at pH 4 and neutral at pH 7. The net charge of the protein is therefore 55 units more positive at pH 4 than at pH 7. These residues are distributed along the entire length of the molecule. Using these assignments as input, we performed independent 10-ns MD simulations at each pH. Backbone C*α* RMSD analysis showed an initial relaxation phase during the first 5 ns in both simulations, with a modest late-stage drift in the pH 4 trajectory; the final 500 frames (5–10 ns) were therefore used for all subsequent analyses (Figure S2C).

We first used the pH 7 MD-relaxed model to resolve the architecture of the C-terminal rod. Structure-based secondary-structure assignment (DSSP/P-SEA) reproduced the two-region partition suggested by the sequence-based predictions (Figure S1), with residues 198–747 forming a predominantly *α*-helical (∼75% helix), markedly elongated rod (≈270 Å; Figure S3A). Knobs-into-holes analysis with SOCKET2 detected a single antiparallel three-helix coiled-coil bundle (helices ∼404–437, 448–475 and 483–510; knobs-into-holes core ∼408–425) and no canonical knobs-into-holes packing elsewhere in the rod (Figure S3B), matching the single coiled-coil locus predicted from sequence (Figure S1B). The C*α*–C*α* contact map of the rod instead showed a tandem series of ∼6 antiparallel helical modules (the characteristic diamond pattern; Figure S3C), indicating an antiparallel helical repeat rather than a parallel coiled-coil arrangement. Together, these analyses define the C-terminal region as an antiparallel *α*-helical, spectrin-repeat-like solenoid that contains one localized coiled-coil.

### 3.3. Profile- and structure-based homology searches

We then asked whether this architecture has recognizable counterparts elsewhere. Profile-based searching with HHpred and structural searching with Foldseek, the latter using the pH 7 MD-relaxed model as the query, returned only weak hits, and these were confined to the C-terminal region; the N-terminal module returned no informative matches. The highest-scoring hits were to elongated *α*-helical proteins, a trypanosomatid microtubule-associated protein (HHpred) and a dystrophin-like protein from a chigger mite (*Leptotrombidium deliense*; Foldseek), aligning over residues 198–720, the same region defined above as the spectrin-repeat-like rod (Tables S1 and S2).

### 3.4. Structural analysis of the 89-kDa periplasmic protein at pH 4 and pH 7

We next compared the equilibrated conformations obtained at the two pH values. Rigid-body domain analysis using DynDom partitioned the protein into an N-terminal core (Domain 1, residues 48–286/299–395), an N-terminal arm (Domain 2, residues 6–30) and a C-terminal rod (Domain 3, residues 287–298/396–747). Two hinge regions were identified: hinge-1 (bending residues 30–48, centered on Pro43) and hinge-2 (bending residues 286–302 and 395–396, centered on Gly291) (Figure 2A).

**Figure 2.**
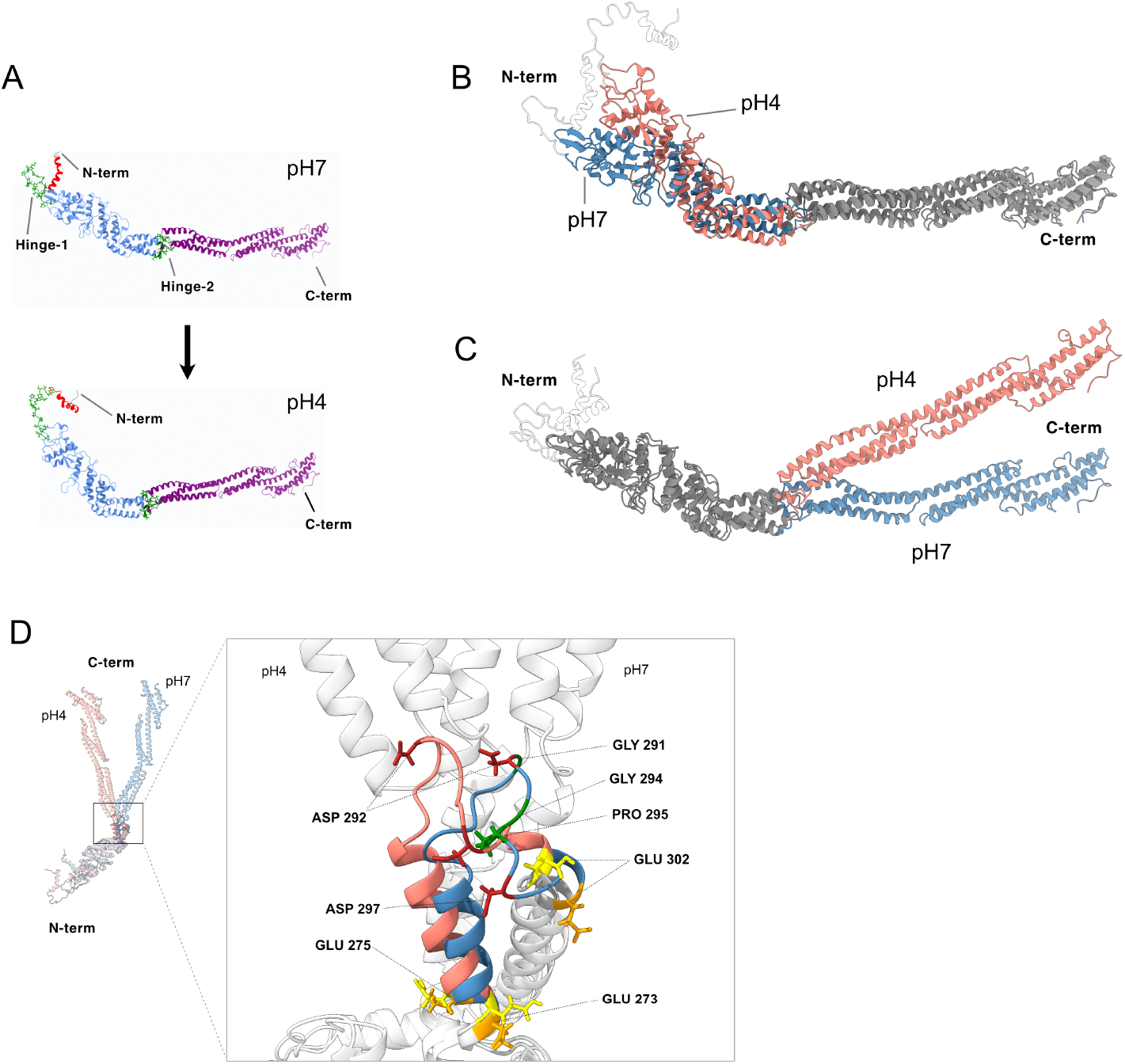
pH-dependent conformational change of the 89-kDa periplasmic invasion-tip protein. (**A**) Ribbon representations of the equilibrated MD structures at pH 7 (top) and pH 4 (bottom), colored according to DynDom domain assignments. Domain 1 (N-terminal core, residues 48–286 and 299–395; blue), Domain 2 (N-terminal arm, residues 6–30; red) and Domain 3 (C-terminal rod, residues 287–298 and 396–747; purple) are shown. Hinge-1 (residues 30–48) and hinge-2 (residues 286–302 and 395–396) are highlighted as green sticks. Unassigned terminal residues are shown in gray. The arrow indicates the pH 7 → pH 4 transition modeled in this study. (**B, C**) Superposition of the equilibrated pH 7 (blue) and pH 4 (salmon) structures on the C-terminal rod (C*α* atoms of residues 396–747; RMSD 3.4 Å over 352 atoms) (**B**) or on the N-terminal core (C*α* atoms of residues 48–286/299–395; RMSD 3.1 Å over 336 atoms) (**C**). In each panel, the region used as the superposition reference is shown in gray. Residues 1–47, which fall within the low-confidence N-terminal region (Figure S2A), are shown as semi-transparent and are not interpreted here. (**D**) Close-up of the hinge-2 region, comparing the pH 7 (blue) and pH 4 (salmon) conformations superposed on the N-terminal core (DynDom Domain 1, residues 48–286/299–395). Left (inset): full-length superposition with the N-and C-termini indicated; the boxed area is enlarged at right. Right: residues 273–302 are colored by state (pH 7, blue; pH 4, salmon) and the remainder is light gray. Side chains are shown for the Gly/Pro hinge residues (Gly291, Gly294, Pro295; green), the flanking glutamates Glu273, Glu275 and Glu302 (charged at pH 7, orange; protonated at pH 4, yellow), and Asp292 and Asp297 (firebrick). Protonation states are PROPKA/PDB2PQR assignments used as simulation inputs.

To characterize this rearrangement further, we superposed the two structures on each of the two large domains in turn. Superposition on the C-terminal rod gave a C*α* RMSD of 3.4 Å over 352 atoms (Figure 2B), and superposition on the N-terminal core gave 3.1 Å over 336 atoms (Figure 2C). In each case the domain used as the reference was closely superimposed, while the remainder of the molecule was displaced. The corresponding per-residue displacement profiles are approximately mirror images of one another, the displacement progressively increasing with distance from the domain junction in both reference frames (Figure S4A). Both domains therefore retain essentially the same internal structure at the two pH values, and the difference between the two states arises predominantly from a change in their relative orientation about the hinge-2 region that connects them, which DynDom quantified as a rigid-body rotation of approximately 35° (Figure 2A). The position of the N-terminal arm is poorly determined in the model (Figure S2A), so its apparent displacement is not interpreted here. Although the precise trajectory of the N-terminal arm cannot be inferred from the present model, exposure of the N-terminal region beyond the bacterial outer membrane during infection has been demonstrated experimentally [14]. A morph animation interpolating between the two models is provided as Video S1; the interpolation is geometric and does not represent a simulated transition pathway. Per-residue RMSF (Figure S4B) showed that hinge-1 and the N-terminal arm substantially fluctuate at both pH values, which is consistent with the low prediction confidence in this region. Both the hinge-2 loop (residues 286–302) and the core–rod junction and proximal rod (residues ∼390–520) showed lower RMSF at pH 4 than at pH 7, with no consistent trend further along the rod (residues >520). Rather than reflecting increased disorder, this suggests that the pH 4 conformation is comparatively well ordered at the hinge-2 loop and along the proximal rod.

We next examined the hinge-2 region in structural detail (Figure 2D). The divergent segment (residues 273–302) contains a glycine/proline-rich loop (Gly291, Gly294, Pro295), flanked by three glutamates (Glu273, Glu275, Glu302) that are predicted to be neutral at pH 4, whereas two aspartates within the same loop (Asp292, Asp297) remain charged at both pH values. DynDom located hinge-2 from the two sets of coordinates alone, without reference to sequence. This geometrically defined hinge coinciding with a glycine/proline-rich loop means that two independent criteria (backbone geometry and amino-acid composition) identify the same segment as the flexible connection between the N-terminal core and the C-terminal rod. Within the loop itself, the conformational change is small: superposed on residues 286–302 alone, the pH 7 and pH 4 models differ by 1.8 Å C*α* RMSD, and the crossing angle between the helices flanking the loop changes by ∼5°. None of the three flanking glutamates forms a salt bridge with a basic residue at either pH.

The connection is not a single joint. DynDom assigns residues 287–298 to the C-terminal rod (Domain 3) even though they lie within the core in sequence, and in the model this segment packs directly against the rod (closest heavy-atom distance 2.8 Å at pH 7 and 2.7 Å at pH 4). The core and the rod are therefore joined at more than one point: the chain passes between them at 286/287, 298/299 and 395/396, rather than through a single linker. The sequence-level features, structural homology, architecture and DynDom domain assignments described above are summarized on a common residue axis in Figure 3.

**Figure 3.**
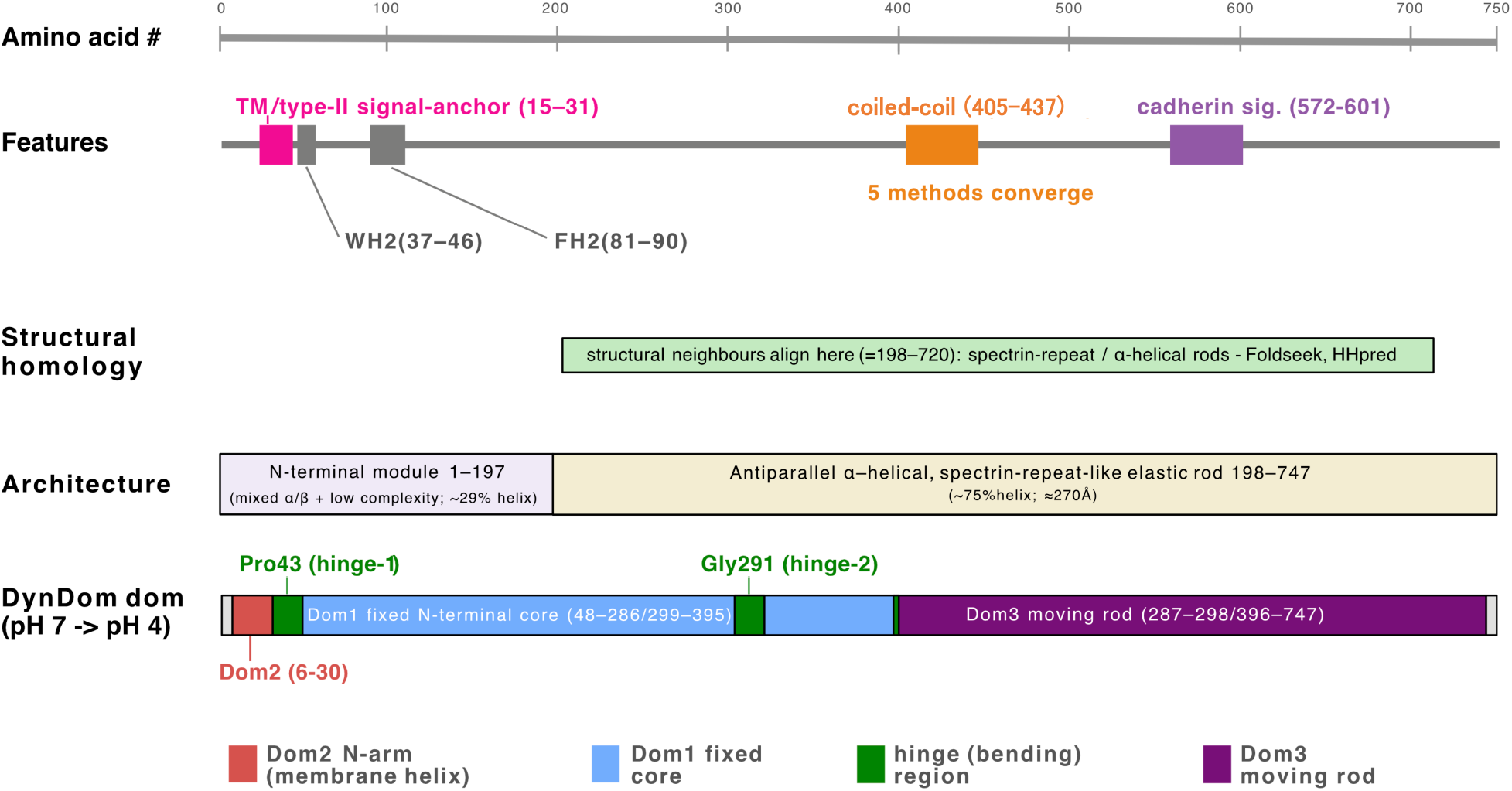
Integrated domain/feature map of the 89-kDa protein, summarizing the homology, primary-to-tertiary structure, and pH-dependent conformational analyses. Tracks are ordered top to bottom from earliest to most recent evidence. Features: sequence-level motifs, including a transmembrane/type-II signal-anchor helix (15–31; TMHMM and PSORT II/ALOM), the two putative WH2-like and FH2-like actin-binding motifs (37–46 and 81–90, respectively) and a cadherin signature (572–601) originally proposed from sequence [14], and a single coiled-coil (405–437) on which five independent methods converge (Figure S1B). Structural homology: the residue range (≈198–720) over which structural neighbors (spectrin-repeat / *α*-helical rods) align in Foldseek and HHpred searches. Architecture: secondary-structure assignment partitions the protein into an N-terminal module (1–197; mixed *α*/β with low-complexity segments, ∼29% helix) and a C-terminal antiparallel *α*-helical, spectrin-repeat-like elastic rod (198–747; ∼75% helix, ≈270 Å) (Figures S1A and S3A). DynDom domains: rigid-body decomposition of the pH 7 → pH 4 conformational change into a fixed N-terminal core (Domain 1, residues 48–286/299–395; blue), a mobile N-terminal arm carrying the membrane helix (Domain 2, residues 6–30; red) and a mobile C-terminal rod (Domain 3, residues 287–298/396–747; purple; rotation 35°), hinged at Pro43 (hinge-1) and Gly291 (hinge-2; green). Dom1–Dom3, Domains 1–3.

## 4. Discussion

Sequence-based homology searches identified confident orthologues only within the genus *Holospora*, with no hits to proteins from other taxa (see Results). This suggests that the 89-kDa protein has either diverged extensively or evolved largely without recognizable homologues outside this genus. This lack of similarity was not uniform along the sequence: weak but detectable hits were recovered for the C-terminal region, whereas the N-terminal module returned no informative matches (Tables S1 and S2). If a bona fide N-terminal homologue is eventually identified in other organisms, it may help to clarify the evolutionary origin of this module. Conversely, if it genuinely remains absent from other lineages, this would be consistent with either an origin unique to *Holospora* or, more speculatively, an evolutionary history in which the N- and C-terminal modules were assembled from sequences of separate origin.

The two C-terminal hits noted above are individually weak, and they should be treated with caution. However, both point toward proteins that act as cytoskeletal or membrane-cortex scaffolds through interaction with partner proteins. Dystrophin in particular anchors the actin cytoskeleton to the plasma membrane via a spectrin-repeat rod. Combined with our finding that the C-terminal region of the 89-kDa protein is itself an antiparallel, spectrin-repeat-like solenoid (Figure 3 and Figure S3), this raises the possibility that the protein does not act alone, but rather assembles with other, as yet unidentified proteins, into a structural complex beneath the *H. obtusa* outer membrane. A further observation bears on this possibility. In addition to the 89-kDa protein, a 76-kDa protein has been detected in micro-dissected invasion-tips of the *H. obtusa* IF cell, and immunofluorescence indicates that it is confined to the invasion-tip (M. Fujishima, unpublished data); its gene has not yet been identified. We have no direct evidence of a physical interaction between the two proteins, but the presence of a second, invasion-tip-specific protein in the same compartment would be consistent with the 89-kDa protein functioning within a complex. If such a complex exists, it could contribute to retaining the C-terminal rod within the periplasmic space while the N-terminal segment becomes exposed at the bacterial surface.

The MD and DynDom analyses indicate that the difference between the pH 7 and pH 4 models is a rigid-body reorientation of the N-terminal core and the C-terminal rod about hinge-2, rather than a deformation of either domain (Figures 2B and 2C). Two features of this arrangement are worth noting. First, the hinge identified from backbone geometry alone coincides with a glycine/proline-rich loop, so that geometry and sequence composition independently point to the same segment as the flexible connection between the two domains (Figure 2D). Second, the core and the rod are joined at two points rather than one: DynDom assigns residues 287–298 to the rod domain although they lie within the core in sequence, and this segment packs directly against the rod in the model. The rod is ∼270 Å long, so a small change at this junction is expressed as a displacement of up to ∼80 Å at its distal end (Figure S4A).

We were unable to identify the interaction through which protonation acts at this junction. None of the three glutamates flanking the hinge-2 loop forms a salt bridge at either pH, and the local rearrangement within the loop is small (see Results). The pH dependence of the reorientation may therefore reflect the distributed change in charge across the molecule as a whole (55 residues; Figure S2B) rather than the rupture or formation of a specific ion pair. Distinguishing between these possibilities will require experiment.

The biological significance of this predicted conformational change lies in its potential connection to the earliest experimentally defined step of *H. obtusa* infection. Previous cell-biological studies have shown that acidification of the host DV is required for bacterial escape [20] and that, during this period, the N-terminal region of the 89-kDa invasion-tip protein becomes exposed beyond the bacterial outer membrane [14]. The pH-dependent reorientation predicted here therefore provides a possible structural link between an environmental cue generated by the host DV and activation of the bacterial invasion apparatus (Figure 4). pH-triggered conformational switches are well established in other systems, notably in viral fusion glycoproteins, in which acidification promotes large-scale structural reorganization [47]. Although the 89-kDa protein is unrelated to these proteins in sequence and fold, this precedent illustrates how acidification can gate a large-scale conformational reorientation. The pH modeled here corresponds to the transient acidification of the DV-I stage, from which escape occurs, and to the condition (pH 4 for 15 min) shown experimentally to induce transformation of the infectious form into the activated form [19]; the more acidic DV-II stage (pH < 3) blocks escape [20].

**Figure 4.**
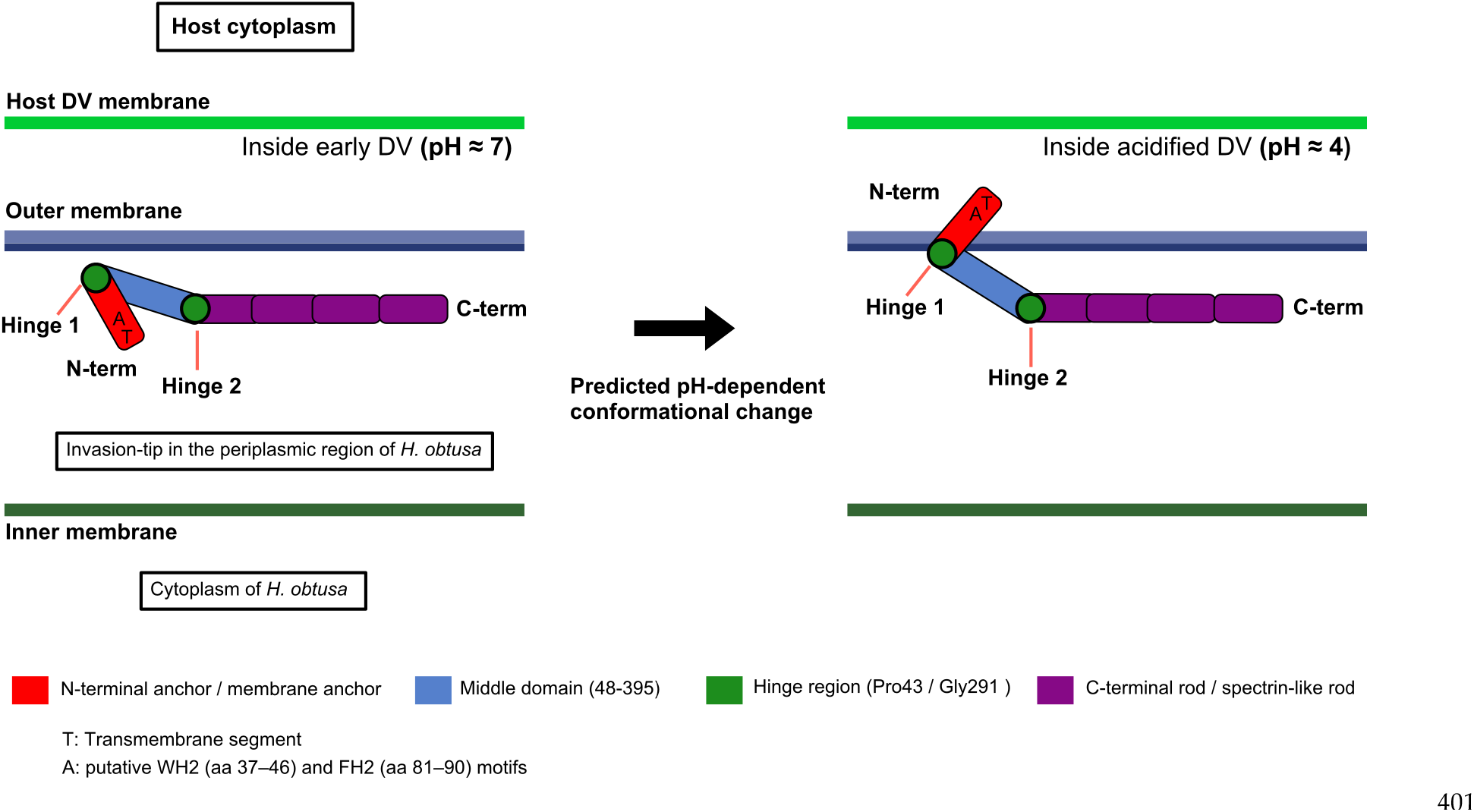
Proposed model of pH-dependent conformational change in the 89-kDa periplasmic invasion-tip protein during escape of the infectious form of *H. obtusa* from the host digestive vacuole (DV). Schematic cross-sections show, from top to bottom, the host cytoplasm, the host DV membrane, the outer and inner membranes of *H. obtusa*, and the intervening periplasmic space, where the invasion-tip protein resides. Left: inside the early, near-neutral DV (pH ≈ 7), the protein adopts an overall extended conformation: the N-terminal membrane anchor (red; T, transmembrane segment; A, putative WH2/FH2-like actin-binding motifs, residues 37–46 and 81–90) lies close to the outer membrane, while the middle domain (blue, residues 48–395) and the C-terminal spectrin-like rod (purple) extend away within the periplasmic space, held in this configuration by hinge-1 (Pro43) and hinge-2 (Gly291; green). Right: upon acidification of the DV (pH ≈ 4), the molecule adopts a more compact, curved overall conformation. In the model, this is accompanied by a reorientation of the N-terminal anchor about hinge-1. The conformation of this segment is poorly determined (Figure S2A), so the schematic shows its repositioning qualitatively rather than to scale.

Per-residue RMSF analysis showed that both the hinge-2 loop and the core–rod junction are more ordered at pH 4 (Figure S4B), suggesting that the pH 4 conformation may represent a discrete state rather than a transiently strained intermediate. Host actin filaments associate with the exposed N-terminal segment following vacuolar escape [16,21]. Actin engagement might therefore restrain the N-terminal arm after entry into the near-neutral host cytoplasm, making an intrinsically reversible con-formational switch functionally unidirectional in vivo. This possibility remains speculative, particularly because the position of the N-terminal arm is poorly determined in the present model (Figure S2A).

Several limitations should be noted in relation to the current study. The model rests entirely on computational prediction and simulation. The three-dimensional structure was generated by AlphaFold2 without an experimental template, and protonation states were assigned computationally (PDB2PQR/PROPKA) rather than measured. The residues identified as titratable and their assigned pKa values may not correspond exactly to their behavior in the native protein, particularly in atypical local environments. Each pH condition was sampled from a single 10-ns trajectory, so we cannot distinguish the differences reported here from the variation that would be expected between independent simulations of the same system. Replicate simulations and longer sampling would thus be required to establish that the reorientation is reproducible. The two states were compared as static endpoints rather than through free-energy or pathway analysis, and the observed difference was not shown to be caused by protonation. The results should therefore be regarded as a structural prediction and a testable hypothesis rather than a demonstrated mechanism. Direct experimental structure determination by X-ray crystallography or cryo-EM at both pH values will be required to test the conformational states proposed here. Finally, subsequent passage of the protein across the host macronuclear envelope appears, from available evidence, to be a separate process that does not obviously depend on the change described here, and is not addressed by the present model.

Experimental testing of this model is constrained by the biology of the organism: *H. obtusa* is an obligate intracellular symbiont that cannot be cultured outside *Paramecium*, and no system for its genetic manipulation is currently available, so direct mutational analysis in the bacterium is not presently feasible. A more accessible route would be to examine the recombinant protein, or fragments of it, in vitro. Whether its overall shape changes between pH 7 and pH 4 could be tested by circular dichroism, size-exclusion chromatography or hydrogen–deuterium exchange mass spectrometry. The contribution of the hinge-2 residues (Glu273, Glu275, Glu302; Gly291, Gly294, Pro295) could be assessed in the same assays using site-directed mutants.

## 5. Conclusions

The 89-kDa invasion-tip protein of *H. obtusa* is predicted to consist of an N-terminal membrane-anchoring module and a long C-terminal antiparallel, spectrin-repeat-like helical solenoid containing a localized coiled-coil. Molecular dynamics simulations at pH 7 and pH 4 revealed a difference in the relative orientation of the N-terminal core and C-terminal rod, described as an approximately 35° rigid-body rotation about a hinge centered on Gly291. Previous cell-biological experiments showed that, after *H. obtusa* IF cell is taken up into a host DV, the N-terminal region of the 89-kDa protein becomes exposed beyond the bacterial outer membrane during vacuolar acidification [14]. The pH-dependent conformational difference predicted in the present study is consistent with this experimentally observed N-terminal exposure and provides a possible structural basis for this early step in the infection process. Although the present computational analyses do not establish that protonation causes this conformational difference, they provide a testable structural hypothesis linking acidification of the host DV to activation of the bacterial invasion apparatus. Experimental structural analyses will be required to determine whether this predicted pH-dependent reorientation occurs during *H. obtusa* infection.

## Supporting information

File S1: Predicted structure at pH 7

File S2: Predicted structure at pH 4

Video S1: Structural morph animation

## Supplementary Materials: Figure S1

Sequence-based secondary-structure, coiled-coil and transmembrane prediction. **Figure S2:** AlphaFold2 model, protonation state and molecular-dynamics setup. **Figure S3:** Structure-based analysis of the C-terminal rod on the molecular-dynamics model. **Figure S4:** Displacement and fluctuation analysis of the pH 7 and pH 4 MD models. **Table S1:** HHpred search of the query BAE80122.1 (89-kDa periplasmic protein, *Holospora obtusa*) against the PDB (job 6845740). **Table S2:** Foldseek 3Di/AA structural search (no iterative search) of the MD-refined model of the query (BAE80122.1, 89-kDa periplasmic protein, *Holospora obtusa*; 750 aa) against AFDB UniProt50 and MGNIFY_ESM30, with no taxonomic filter. **File S1:** Equilibrated structure of the 89-kDa protein at pH 7 (final frame of the 10-ns MD trajectory; PDB format). **File S2:** Equilibrated structure of the 89-kDa protein at pH 4 (final frame of the 10-ns MD trajectory; PDB format). **Video S1:** Morph animation of the predicted pH-dependent conformational change of the 89-kDa periplasmic invasion-tip protein (pH 7 → pH 4), generated with ChimeraX. Coloring follows Figure 2A: N-terminal core (Domain 1, blue), N-terminal arm (domain 2, red), C-terminal rod (Domain 3, purple), hinge regions (green). The interpolation is geometric and does not represent a simulated transition pathway.

## Author Contributions

Conceptualization, M.F. and Y.T.; methodology, Y.T. and M.F.; software, Y.T.; validation, T.M., T.N. and Y.S.; investigation, Y.T.; resources, M.F.; data curation, Y.T.; writing—original draft preparation, M.F. and Y.T.; writing—review and editing, M.F., Y.T., T.M., T.N. and Y.S.; visualization, M.F. and Y.T.; supervision, M.F.; project administration, M.F. and Y.T. All authors have read and agreed to the published version of the manuscript.

## Institutional Review Board Statement

Not applicable.

## Informed Consent Statement

Not applicable.

## Data Availability Statement

The equilibrated pH 7 and pH 4 structures analyzed in this study are provided as Supplementary Materials. The full molecular-dynamics trajectories will be made available by the authors on request.

## Acknowledgments

We acknowledge proofreading and editing by Benjamin Phillis, a Board-Certified Editor in the Life Sciences (BELS), at Wakayama Medical University.

## Abbreviations

The following abbreviations are used in this manuscript:

RF: Reproductive form
IF: Infectious form
DV: Digestive vacuole
AF: Activated form

