## Supplementary Figures S1-S4 and Tables S1-S2 for "Predicted structure of the 89-kDa invasion-tip protein of *Holospora obtusa*, an endonuclear symbiont of *Paramecium caudatum*, suggests a pH-dependent conformational change"

### Figure S1

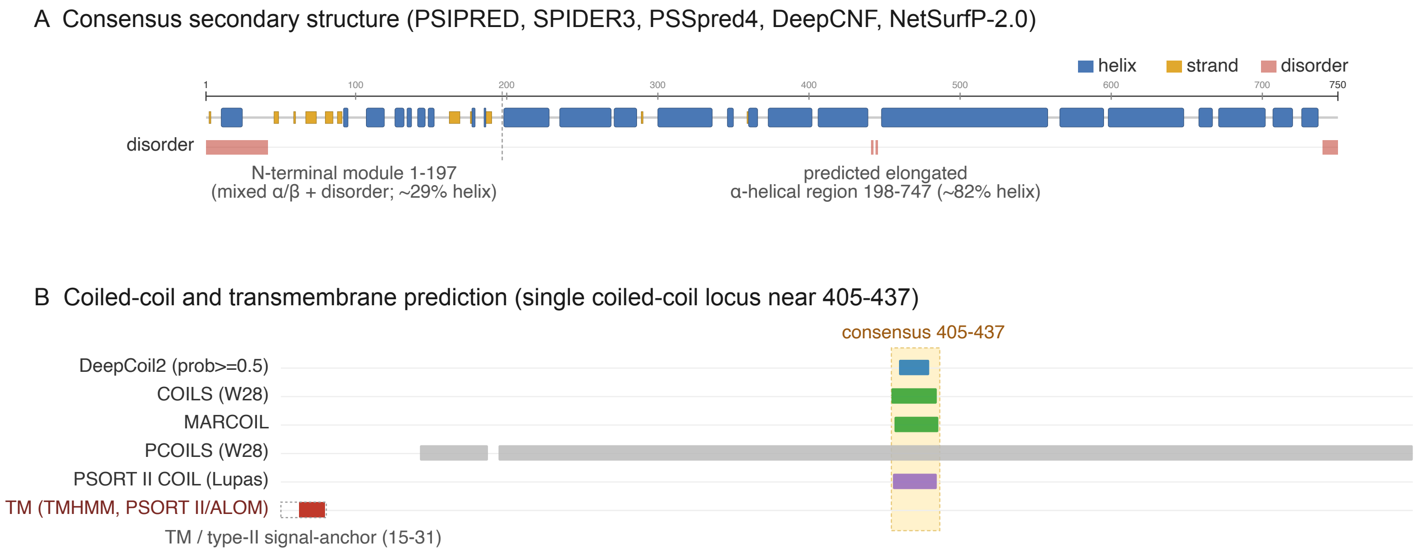


**Figure S1:** Sequence-based secondary-structure, coiled-coil and transmembrane prediction.
All predictions were computed from the 750-residue amino-acid sequence of the 89-kDa invasion-tip protein using the MPI Bioinformatics Toolkit, DeepCoil2 and PSORT II. (A) Consensus secondary structure derived from five independent predictors (PSIPRED, SPIDER3, PSSpred4, DeepCNF and NetSurfP-2.0), mapped along the sequence (residues 1–750). Boxes denote helix (blue), strand (yellow) and disorder (pink). The N-terminal module (residues 1–197) is a mixed α/β region with local disorder (~29 % helix), whereas residues 198–747 are predicted as a largely continuous, elongated α-helical region (~82 % helix). (B) Coiled-coil and transmembrane predictions shown as tracks along the same axis: coiled-coil propensity from DeepCoil2 (probability ≥ 0.5), COILS (window 28), MARCOIL, PCOILS (window 28) and the PSORT II COIL (Lupas) method; and the predicted N-terminal transmembrane / type-II signal-anchor (residues 15–31; TMHMM and PSORT II/ALOM). Apart from the PCOILS outlier, the methods converge on a single coiled-coil locus at residues ~405–437 (consensus, shaded). PCOILS (window 28) over-predicts coiled coil across continuous α-helical regions and is the only discordant track; this locus is validated on the structure in Fig. S3.

### Figure S2

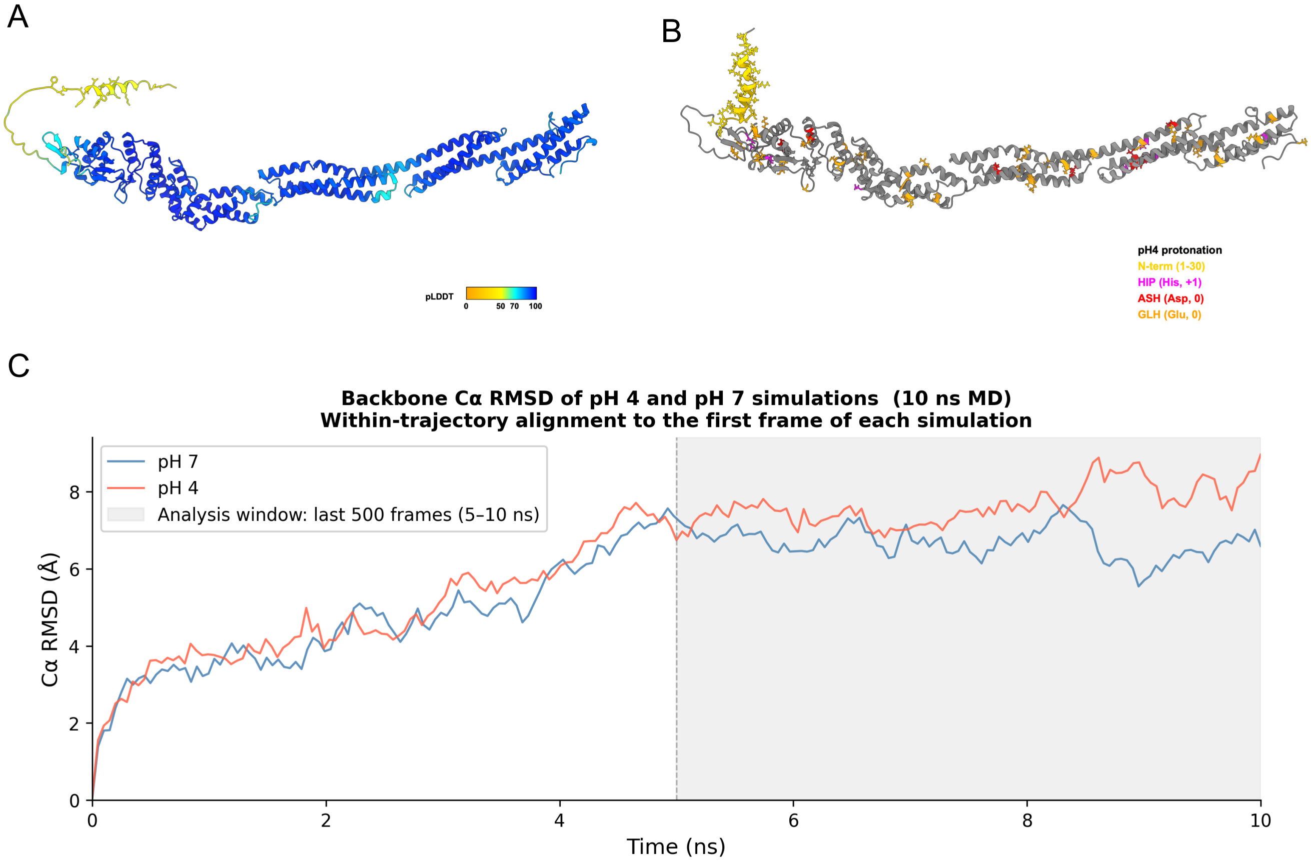


**Figure S2:** AlphaFold2 model, protonation state and molecular-dynamics setup. (A) AlphaFold2-predicted structure of the 89-kDa invasion-tip protein, colored by per-residue confidence score (pLDDT). The N-terminal region (residues 1–30) exhibits low pLDDT values (< 50, yellow). (B) Distribution of pH-sensitive residues mapped onto the 89-kDa protein structure (final frame of 10 ns MD simulation at pH 7; gray ribbon). Residues predicted by PDB2PQR to change protonation state between pH 4 and pH 7 are shown as sticks: His residues protonated at pH 4 (HIP; n = 7, magenta), Asp residues neutral at pH 4 (ASH; n = 8, red), and Glu residues neutral at pH 4 (GLH; n = 39, orange). The N-terminal region (residues 1–30) is also shown as sticks (gold). The net charge difference between pH 4 and pH 7 is 55 e. (C) Backbone Cα RMSD time course of pH 7 and pH 4 MD simulations. RMSD was calculated for backbone Cα atoms after structural alignment to the first frame of each trajectory. Both simulations showed an initial relaxation phase during the first 5 ns, followed by late-trajectory behavior within an approximately 6–9 Å range. The shaded region indicates the late-trajectory analysis window corresponding to the last 500 frames of the original trajectory files, equivalent to 5–10 ns. The relatively large RMSD values are consistent with the substantial displacement of the N-terminal arm and interdomain rearrangement detected by DynDom analysis.

### Figure S3

A Secondary structure on the model (DSSP / Cα P-SEA)

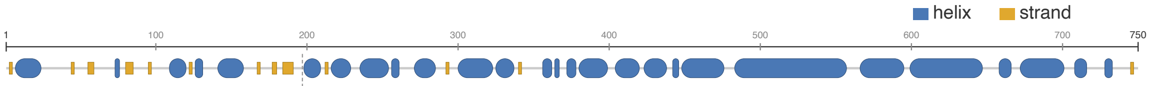


N-terminal module 1-197 (~29% helix) antiparallel α-helical, spectrin-repeat-like rod 198-747 (~75% helix; ≈270 Å)

B SOCKET2 coiled-coil assignment (cutoff 7.0 Å)

| | |
|---|---|
| Total α-helical residues | 471 |
| Coiled coils detected | 1 (antiparallel, 3-stranded) |
| Helices in bundle | H15 404-437 · H17 448-475 · H18 483-510 |
| Knobs-into-holes core | ≈ residues 408-425 |
| Length (max / mean) | 19 / 15 residues |
| Heptad register | clean (0 non-canonical interrupts) |
| Rest of rod (198-747) | no canonical knobs-into-holes |

C Cα contact map of the rod (198-720)

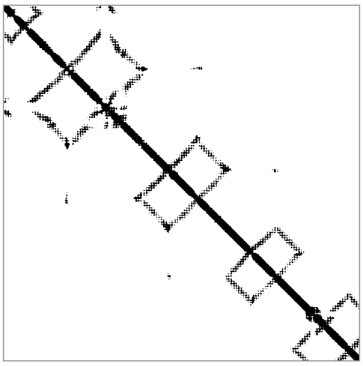

~6 tandem antiparallel modules (diamond pattern)
contacts < 12 Å; axes = residues 198-720

**Figure S3:** Structure-based analysis of the C-terminal rod on the molecular-dynamics model. All panels were computed from the final frame of the pH 7 MD trajectory of the 89-kDa invasion-tip protein. (A) Secondary structure assigned on the model by DSSP and Cα P-SEA, mapped along the sequence (residues 1–750); helix (blue), strand (yellow). The N-terminal module (residues 1–197) is ~29 % helical, while residues 198–747 form an antiparallel α-helical, spectrin-repeat-like rod (~75 % helix, ≈270 Å long). (B) Knobs-into-holes packing assigned by SOCKET2 (cutoff 7.0 Å). Of 471 α-helical residues, a single canonical coiled coil is detected: an antiparallel, three-stranded bundle formed by helices H15 (404–437), H17 (448–475) and H18 (483–510), with a knobs-into-holes core at ~residues 408–425 (segment length 19/15 residues, max/mean) and a clean heptad register (no non-canonical interruptions). The remainder of the rod (residues 198–747) contains no canonical knobs-into-holes packing. (C) Cα–Cα contact map of the rod (residues 198–720; contacts < 12 Å). The off-diagonal "diamonds" reveal ~6 tandem antiparallel modules stacked along the rod axis.

### Figure S4

A

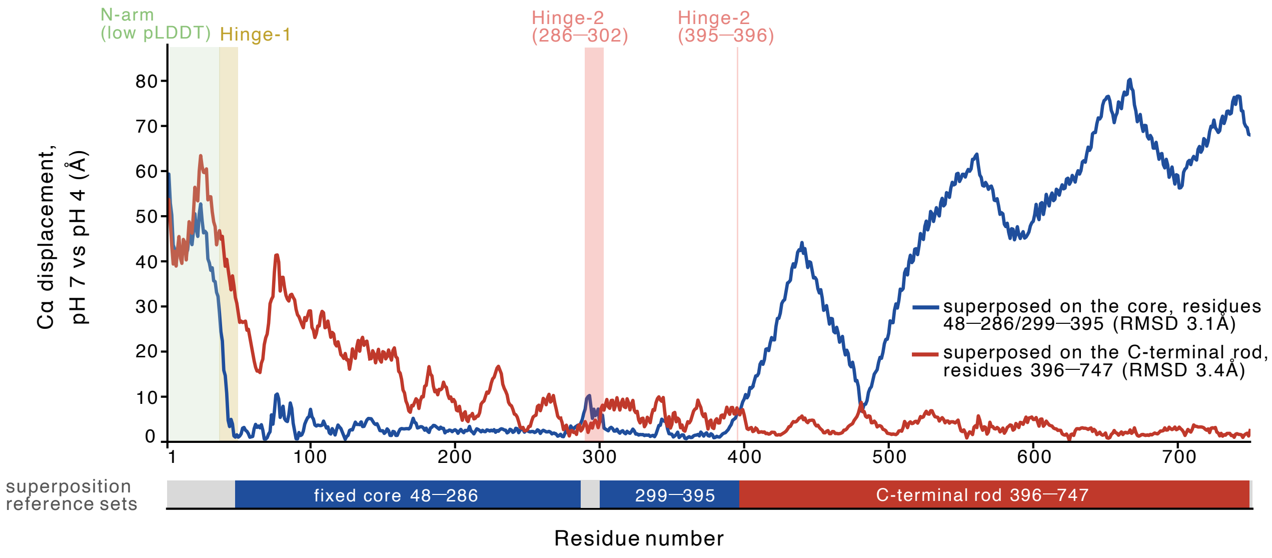


B

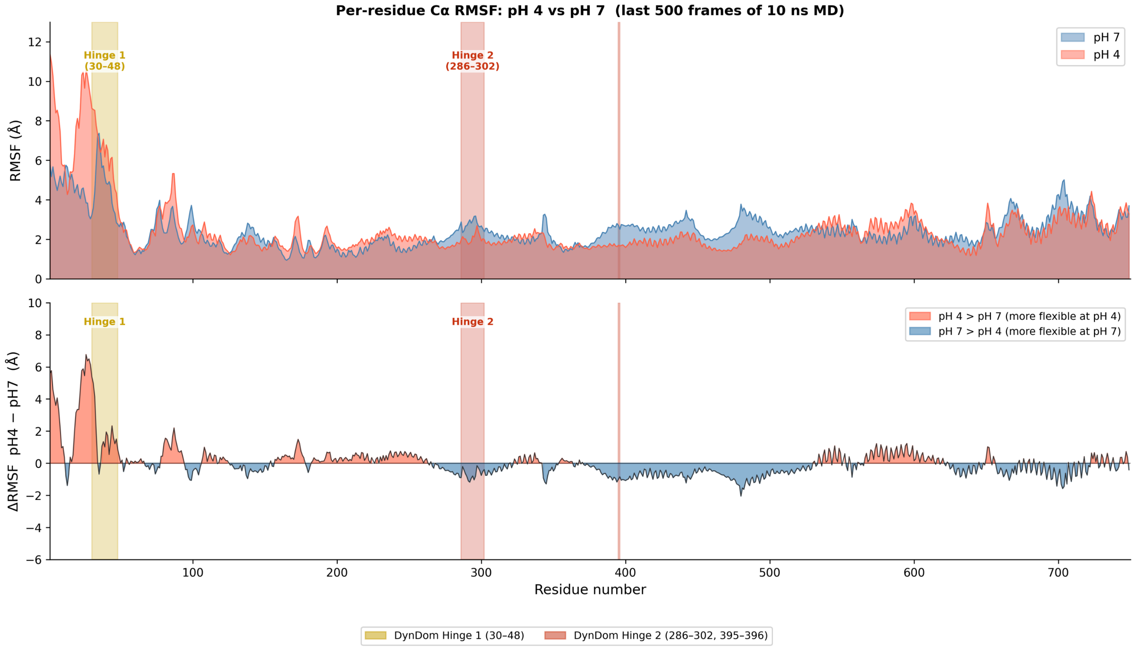


**Figure S4:** Displacement and fluctuation analysis of the pH 7 and pH 4 MD models. (A) Per-residue Cα displacement between the pH 7 and pH 4 final-frame structures of the 89-kDa invasion-tip protein, computed twice using different superposition reference sets: the N-terminal core domain (residues 48–286/299–395; blue; RMSD 3.1 Å) or the C-terminal rod (residues 396–747; red; RMSD 3.4 Å). The bar below indicates the reference sets. Shaded bands indicate DynDom-defined hinge regions: hinge-1 (residues 30–48) and hinge-2 (residues 286–302 and 395–396). (B) Per-residue Cα root-mean-square fluctuation (RMSF) of the 89-kDa protein at pH 7 (blue) and pH 4 (red), calculated from the Cα B-factors of the last 500 frames of the 10-ns MD trajectory at each pH. Top: absolute RMSF for each condition. Bottom: difference (ΔRMSF = pH 4 − pH 7); red, more flexible at pH 4; blue, more flexible at pH 7. Shaded bands as in (A).

#### Table S1

| Nr | Hit | Name | Structural class | Probability | E-value | Score | SS | Aligned cols | Target Length |
|---|---|---|---|---|---|---|---|---|---|
| 1 | 9E5C_1c | CMF5; flagella, microtubule, MOTOR PROTEIN; HET: MG, GTP, ATP, ADP, GDP, ZN; 3.2A (Trypanosoma brucei TREU927) | Microtubule / axoneme | 96.11 | 5.4 | 46.6 | 26.2 | 315 | 1340 |
| 2 | 7NRC_A | GCN1; Ribosome, Disome, GCN1, Translation, GAAC, ISR, Rbg2, Gir2; HET: 5CT; 3.9A (Saccharomyces cerevisiae S288C) | HEAT-repeat solenoid | 96.03 | 0.71 | 58.1 | 19.4 | 252 | 1209 |
| 3 | 9E5C_1d | CMF6; flagella, microtubule, MOTOR PROTEIN; HET: MG, GTP, ATP, ADP, GDP, ZN; 3.2A (Trypanosoma brucei TREU927) | Microtubule / axoneme | 95.95 | 5.8 | 45.57 | 27.7 | 329 | 1204 |
| 4 | 9Y6S_9W | Guanine nucleotide-binding protein subunit beta-like protein; flagella, parasite, axoneme, doublet microtubule, STRUCTUR (Leishmania tarentolae) | Microtubule / axoneme | 95.82 | 6.5 | 45.2 | 29.1 | 321 | 1236 |
| 5 | 9Y6S_9X | Guanine nucleotide-binding protein subunit beta-like protein; flagella, parasite, axoneme, doublet microtubule, STRUCTUR (Leishmania tarentolae) | Microtubule / axoneme | 95.74 | 5.5 | 49.71 | 25.1 | 234 | 1593 |
| 6 | 8GLV_Cp | WD_REPEATS_REGION domain-containing protein; cilia, microtubule, dynein, MOTOR PROTEIN; HET: GTP, MG, GDP; 3.1A (Chlamydomonas reinhardtii) | Microtubule / axoneme | 95.45 | 9.9 | 44.73 | 24.4 | 239 | 1316 |
| 7 | 8J07_i4 | Cilia- and flagella-associated protein 57; cilia, microtubule, dynein, STRUCTURAL PROTEIN; HET: GDP, GTP, MG; 4.1A (Homo sapiens) | Microtubule / axoneme | 95.29 | 10 | 44.02 | 25.2 | 329 | 1250 |
| 8 | 9Y6S_9W | Guanine nucleotide-binding protein subunit beta-like protein; flagella, parasite, axoneme, doublet microtubule, STRUCTUR (Leishmania tarentolae) | Microtubule / axoneme | 95.26 | 10 | 43.73 | 24.7 | 330 | 1236 |
| 9 | 8AX9_A | Maltose/maltodextrin-binding periplasmic protein,Apolipoprotein E; Apolipoprotein E, LIPID BINDING PROTEIN; 1.549A (Homo sapiens) | Other / low relevance | 94.57 | 10 | 41.98 | 20.5 | 220 | 675 |
| 10 | 6R1J_D | Uncharacterized protein; Tripartite pore-forming toxin, TOXIN; 1.92A (Aeromonas hydrophila J-1) | Other / low relevance | 94.51 | 9.3 | 39.63 | 21.4 | 209 | 272 |

**Table S1:** HHpred search of the query BAE80122.1 (89-kDa periplasmic protein, *Holospora obtusa*) against the PDB (job 6845740); the top 10 hits ranked by Probability are shown. Metrics: Probability, HHpred's calibrated estimate that the template is a true homolog of the query; E-value, number of chance hits expected at this score or better; Score, raw profile-profile alignment score; SS, the secondary-structure contribution to the Score. Hits are grouped into three structural classes, shown by row color: ■ Microtubule / axoneme (Nr 1, 3-8); ■ HEAT-repeat solenoid (Nr 2); ■ other / low relevance (Nr 9, 10). Most hits lie in the twilight zone (Probability 94-96%, E-value ~0.7-10), so differences in individual rank are not significant (the best, Nr 2 / 7NRC, has E-value 0.71). The convergence on microtubule/axoneme structures may partly reflect the abundance of such cryo-EM complexes in the PDB and should be read as consistent with this fold class rather than as proof of function.

### Table S2

#### AFDB50 (AFDB UniProt50) — top 10 hits

| Target | Description | Scientific Name | Group (relevance) | Prob. | Seq. Id. | E-Value | Score | Query Pos. | Target Pos. |
|---|---|---|---|---|---|---|---|---|---|
| AF-Q2ABW8-F1-model_v6 | 89-kDa periplasmic protein | *Holospora obtusa* | Holospora homolog (true positive) | 1.00 | 99.8 | 4.81e-84 | 3525 | 1–750 (750) | 1–750 (750) |
| AF-A0A2S5RHV0-F1-model_v6 | 89-kDa periplasmic protein | *Holospora curviuscula* | Holospora homolog (true positive) | 1.00 | 95.5 | 3.57e-45 | 1899 | 28–748 (750) | 22–738 (740) |
| AF-A0A0A61J449-F1-model_v6 | Uncharacterized protein | *Holospora undulata* HU1 | Holospora homolog (true positive) | 1.00 | 34.1 | 2.82e-44 | 1763 | 1–743 (750) | 1–747 (750) |
| AF-A0A0A2C0XH1-F1-model_v6 | Uncharacterized protein | *Holospora elegans* E1 | Holospora homolog (true positive) | 1.00 | 32.9 | 1.33e-33 | 1281 | 1–587 (750) | 1–582 (583) |
| AF-A0A0A2C0Z25-F1-model_v6 | Uncharacterized protein | *Holospora elegans* E1 | Holospora homolog (true positive) | 1.00 | 22.2 | 1.43e-27 | 813 | 30–738 (750) | 1–719 (727) |
| AF-A0A813JFD3-F1-model_v6 | Uncharacterized protein (Fragment) | *Polarella glacialis* | Distant fold-level hit (α-helical rod) | 1.00 | 8.5 | 3.66e-3 | 108 | 198–738 (750) | 148–666 (671) |
| AF-A0A7S1R9X9-F1-model_v6 | I/LWEQ domain-containing protein (Fragment) | *Alexandrium catanella* | Distant fold-level hit (α-helical rod) | 1.00 | 8 | 1.48e-2 | 107 | 199–733 (750) | 73–558 (588) |
| AF-A0A443SBB9-F1-model_v6 | Dystrophin-like protein (Fragment) | *Leptotrombidium deliense* | Distant fold-level hit (α-helical rod) | 1.00 | 12.4 | 4.02e-3 | 106 | 104–733 (750) | 19–553 (559) |
| AF-A0A183CZB1-F1-model_v6 | Dystrophin | *Gongylonema pulchrum* | Distant fold-level hit (α-helical rod) | 1.00 | 10.3 | 2.22e-2 | 98 | 198–717 (750) | 17–486 (533) |
| AF-A0A813KJU9-F1-model_v6 | Uncharacterized protein | *Polarella glacialis* | Distant fold-level hit (α-helical rod) | 1.00 | 11.8 | 2.78e-2 | 97 | 198–734 (750) | 250–762 (1017) |

#### MGNIFY_ESM30 — top 10 hits

| Target | Group (relevance) | Prob. | Seq. Id. | E-Value | Score | Query Pos. | Target Pos. |
|---|---|---|---|---|---|---|---|
| MGYP000265513544 | Metagenomic structural analog (unannotated) | 0.99 | 10.1 | 8.03e-2 | 88 | 198–715 (750) | 50–504 (508) |
| MGYP000380261023 | Metagenomic structural analog (unannotated) | 0.99 | 13.7 | 1.50e-1 | 82 | 252–580 (750) | 2–305 (361) |
| MGYP001437905195 | Metagenomic structural analog (unannotated) | 0.99 | 11.2 | 5.36e-2 | 81 | 198–735 (750) | 10–488 (511) |
| MGYP003555838343 | Metagenomic structural analog (unannotated) | 0.98 | 11.9 | 5.86e-2 | 80 | 271–717 (750) | 3–398 (398) |
| MGYP001797285657 | Metagenomic structural analog (unannotated) | 0.98 | 11 | 5.36e-2 | 78 | 197–721 (750) | 17–492 (492) |
| MGYP001501797793 | Metagenomic structural analog (unannotated) | 0.97 | 10 | 2.47e-1 | 75 | 254–612 (750) | 2–324 (325) |
| MGYP002838043246 | Metagenomic structural analog (unannotated) | 0.96 | 16 | 6.86e+0 * | 73 | 391–530 (750) | 1–133 (197) |
| MGYP000879415000 | Metagenomic structural analog (unannotated) | 0.96 | 13.3 | 5.01e+0 * | 73 | 408–643 (750) | 14–239 (250) |
| MGYP003571887057 | Metagenomic structural analog (unannotated) | 0.95 | 12.5 | 2.47e-1 | 72 | 221–719 (750) | 5–451 (459) |
| MGYP000312247625 | Metagenomic structural analog (unannotated) | 0.94 | 12.6 | 1.15e-1 | 71 | 198–673 (750) | 90–506 (510) |

**Table S2:** Foldseek 3Di/AA structural search (no iterative search) of the MD-refined model of the query (BAE80122.1, 89-kDa periplasmic protein, *Holospora obtusa*; 750 aa) against AFDB50 (AFDB UniProt50) and MGNIFY_ESM30, with no taxonomic filter; the top 10 hits per database, ranked by Prob., are shown. Metrics: Prob., Foldseek's estimate that the target is a true structural match (0-1); Seq. Id., percent sequence identity within the structural alignment; E-Value, number of chance matches expected at this score or better; Score, raw Foldseek bit score; Query Pos. / Target Pos., aligned residue ranges (with full length in parentheses). Hits are grouped by relevance, shown by row color: ■ Holospora homolog (true positive) (genus *Holospora*, Seq. Id. 22-100%); ■ Distant fold-level hit (antiparallel α-helical, spectrin-repeat-like rod) (other taxa, Seq. Id. ~8-12%, dystrophin/ILWEQ-type rods); ■ Metagenomic structural analog (unannotated) (MGnify environmental ORFs, no taxonomy/function). * MGNIFY E-Value > 1 (structurally weak). Interpretation. Unlike the earlier search against Swiss-Prot + PDB100 (which lacked *Holospora* and returned spurious human/mouse rods), AFDB50 contains *Holospora* entries and recovers the true homologs: the five top hits are the protein's own orthologs/paralogs in *H. obtusa*, *H. curviuscula*, *H. undulata* and *H. elegans*, confirming the database-composition bias of the previous result. Beyond these, only distant fold-level matches remain (dystrophin / I-LWEQ spectrin-repeat rods at <12% identity), consistent with an elongated α-helical, spectrin-repeat-like rod fold. The Holospora homologs align over the full length (≈1-750), whereas all distant and metagenomic hits align only to ≈residues 198-720, suggesting that the conserved structural module is the C-terminal ~550 residues and that the N-terminal ~1-197 region is Holospora-specific. MGNIFY_ESM30 hits are unannotated metagenomic ORFs with no taxonomy or function and only ~10-16% identity; they add no functional or phylogenetic information, but corroborate that this rod fold is widely distributed among uncultured environmental microbial proteins and is therefore a genuine, conserved architecture rather than a modeling artifact.
